# Evolution of a Plastic Surgery Summer Research Program

**DOI:** 10.1101/2022.05.16.492191

**Authors:** Allyson R. Alfonso, Zoe P. Berman, Gustave K. Diep, Jasmine Lee, Elie P. Ramly, J. Rodrigo Diaz-Siso, Eduardo D. Rodriguez, Piul S. Rabbani

## Abstract

**Background:** Early surgical exposure and research fellowships have been independently shown to influence medical students’ specialty choice, increase academic productivity, and impact residency match. However, to our knowledge there is no published guidance on the implementation of formal plastic surgery summer research programs for first year medical students. We present our institutional experience developing a plastic surgery summer research program over seven years (2013-2020) in an effort to inform program development at other institutions. We hypothesized that this early, formal exposure could spark interest in pursuing research activities throughout medical school and residency.

**Methods:** From 2013 to 2016, a sole basic science research arm existed. In 2017, a clinical research arm was introduced, with several supplemental activities including structured surgical skills sessions. A formalized selection process was instituted in 2014. Participant feedback was analyzed on a yearly basis. Long-term outcomes included continued research commitment, productivity, and residency match.

**Results:** The applicant pool has reached 96 applicants in 2019, with 85% from outside institutions. Acceptance rate reached 7% in 2020. With adherence to a scoring rubric for applicant evaluation, good to excellent interrater reliability was achieved (ICC = 0.75). Long-term outcomes showed that on average per year, 28% of participants continued departmental research activities and 29% returned for dedicated research. Upon finishing medical school, participants had a mean of 6.9±4.0 peer-reviewed publications. 62% of participants matched into a surgical residency program, with 54% in integrated plastic surgery.

**Conclusions:** A research program designed for first year medical students interested in plastic surgery can achieve academic goals. Students are provided with mentorship, networking opportunities, and tools for self-guided learning and career development.

## INTRODUCTION

The continued growth of plastic surgery as an innovative field depends on training the next generation of plastic surgeons. Early surgical exposure has been recognized as the most important factor influencing medical students in choosing a surgical specialty when surveying medical students and residents.^1-3^ In a study evaluating the impact of surgical exposure on medical students’ specialty choice, the implementation of an eight-week surgical research program was shown to result in interest in surgery, with favorable match outcomes.^4^ In addition to assisting career decision-making, the need for early exposure to plastic surgery is further evidenced by medical students’ lack of understanding of the scope of the field, which can negatively impact future referral patterns.^5,6^

A research fellowship with a structured curriculum not only provides medical students with early exposure to plastic surgery and generates interest in the field, but it can also be highly beneficial for residency applicants. The integrated plastic surgery residency track is among the most competitive specialties, with the applicant pool including some of the highest United States Medical Licensing Examination scores and greatest number of Alpha Omega Alpha Honor Medical Society members.^7,8^ A research fellowship provides the opportunity for students to improve their oral presentation experience and publication record, as well as develop a valuable professional network by working closely with academic surgeons and presenting at national meetings.^9^ This supports the ability to obtain high-quality letters of recommendation, which was ranked in a national survey of plastic surgery program directors as one of the most important contributors in selecting residents. Additionally, the contributions of plastic surgery research fellows have historically been important, with published institutional experiences showing significant increase in academic research productivity with the incorporation of formalized research fellowships.^10,11^ Ultimately, applicants who complete a research fellowship demonstrate significantly higher match rates than those who do not.^12^

Currently, to our knowledge there is no published guidance on the implementation of formal plastic surgery summer research programs for first year medical students. In this study, we present our institutional experience developing a plastic surgery summer research program for first year medical students. Through participant feedback, institutional support, and engagement of senior medical students, post-doctoral research fellows, and department faculty, we were able to establish and continuously refine a successful research program.

## METHODS

This study presents an iterative quality enhancement process implementing a plastic surgery summer research program for first year medical students at our institution over seven years (2013-2020). We designed the program to provide a structured learning experience for first year medical students by utilizing a curriculum and learning objectives that affords students the opportunity to harness critical thinking and fundamental research skills, including the ability to identify problems, conceptualize and formulate research questions, design and troubleshoot study approaches, collaborate within a diverse team, and communicate effectively through writing and public speaking. The program is structured to enable students to develop leadership and mentorship expertise, as well as project management skills within a team composed of peers, senior medical students, research fellows, administrators and faculty.

### Initial Program Development

The first iteration of the summer research program in 2013 filled the gap of limited plastic surgery research experience for students, but was unstructured without clearly defined curricular objectives or mentor and mentee roles. Early feedback allowed recognition of an opportunity to more clearly delineate expectations and goals of the program. By evolving an eight-week research curriculum that familiarizes students with scientific literature and development of critical appraisal skills, we hypothesized that students who participated in the program would enter their fields better equipped to incorporate research into their clinical practice. In the short-term, this early, formal exposure could spark interest in pursuing research activities throughout medical school and residency.

Additional focus was placed on maximizing mentor and mentee productivity within an eight-week period. From 2013 to 2016, a sole basic science research arm existed. In 2014, we introduced a formal application process, with established start and end dates. We chose to reduce the number of informal volunteers in favor of an application-based selection process, to better identify committed and qualified students who would derive maximum benefit. The smaller group also allowed mentees to work in closer proximity to research staff, receiving individualized attention and developing self-reliance and independence. Aligned with the mission of the teaching medical institution, projects progressed more steadily.

Over the next four years, the programming team gathered data to present to the Department to support funding avenues and attract a larger applicant pool. This program has since been completely funded by the Department. Each year, we incorporated specific feedback from participants in order to improve communication and tailor management and programming. In 2017, we introduced a clinical research arm with similar goals of applying scientific curiosity to frame research questions, evaluating the primary literature, developing command over a topic, and discussing issues in context. Supplemental activities incorporated with this phase of the program included weekly journal clubs, lab meetings, and surgical skills sessions. Participants attended weekly departmental Grand Rounds and participated in Research Day, which evolved into ultimately presenting their work in front of the Department. The incorporation of this additional programming is also supported by survery data of what trainees desire from their teachers in order to improve their educational experiences. Program expansion allowed for one-to-one mentorship from senior research fellows, with a ratio of two to three students to every faculty member.

### Yearly Planning

Figure 1 delineates the most up-to-date yearly planning timeline. Execution of each category of tasks requires input from multiple groups within our programming team. Success is achieved through the cohesive efforts of financial administrators, pre- and post-doctoral research fellows including residents, senior medical students, and faculty. A central project manager role is necessary for coordination and timely execution of all steps. This organizational model functions as a multi-level tiered training system, which provides a mutual mentorship experience for all involved.

**Figure 1.**
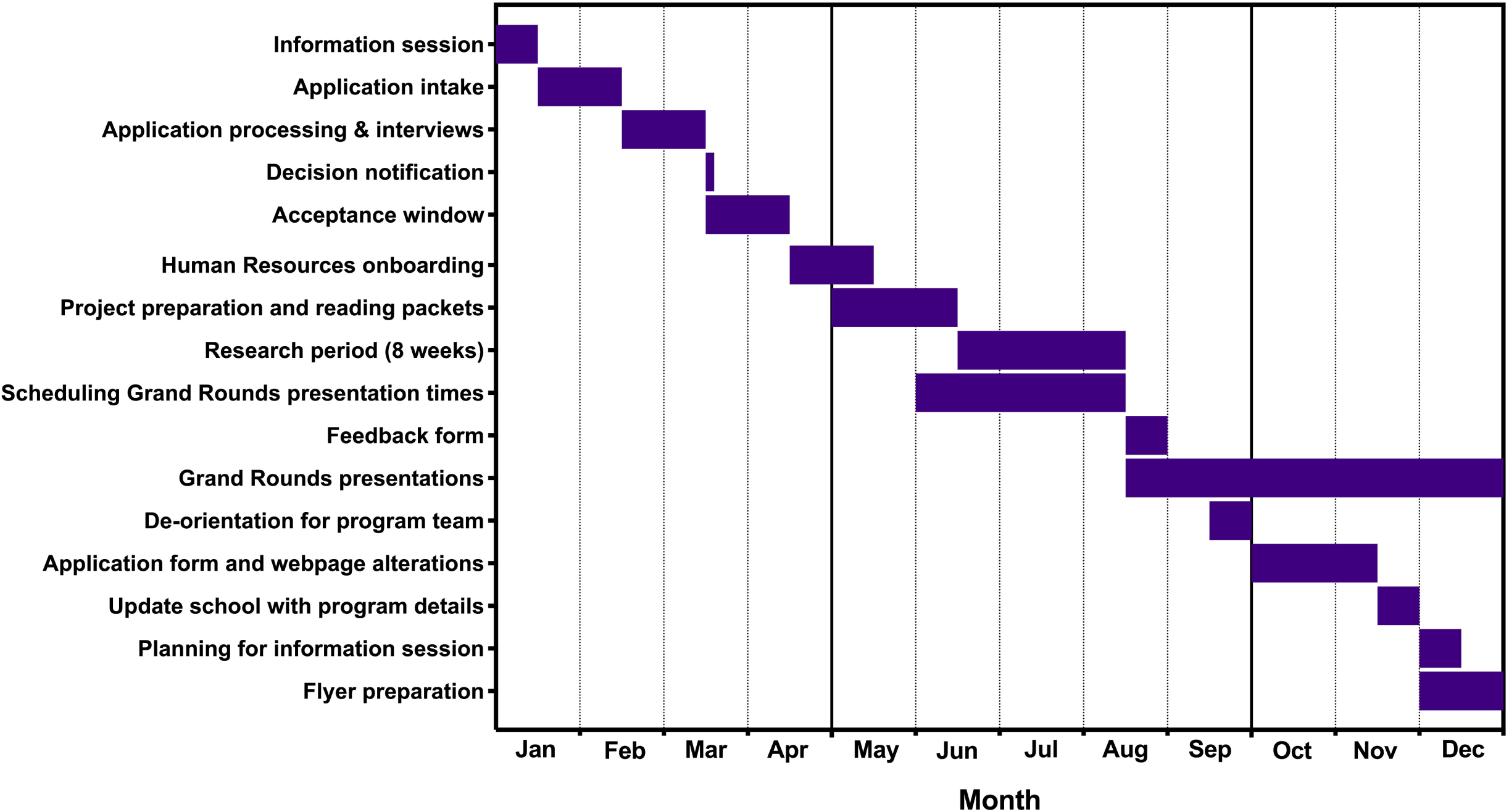
Yearly planning timeline

### Information Session

Recruitment through the online platform^13^ is supplemented by face-to-face efforts. In coordination with the Office of Student Affairs at NYU Grossman School of Medicine, the programming team and program alumni participate in a town hall-style information session followed by a Summer Program Fair. Program alumni serve a critical role as their perspective is highly valued by prospective students. A printed flyer which contains program highlights supplements the information shared with students. Follow-up questions are addressed by team members via email and/or phone.

### Application and Student Selection Process

Presently, only online applications are accepted. **Table 1** demonstrates the application questions. In order to review upwards of 100 applications, we assign each application to two evaluators using a randomized assignment generator.^14^ For the most recent application cycle, we developed a scoring rubric (**Figure 2**) to grade applications according to seven criteria. The rubric was derived from online educational resources and instruction literature.^15,16^ The rubric allows for a focused and standardized assessment of each candidate’s application. Rubric topics also guide collaborative discussion amongst evaluators to promote holistic assessment of each candidate. A heterogeneous panel of evaluators from the research team are responsible for assessing the applicant pool. Most recently, this panel consisted of two senior medical students in pre-doctoral research fellow roles, three residents in full-time post-doctoral research fellow roles, and one basic science faculty member.

**Table 1.**
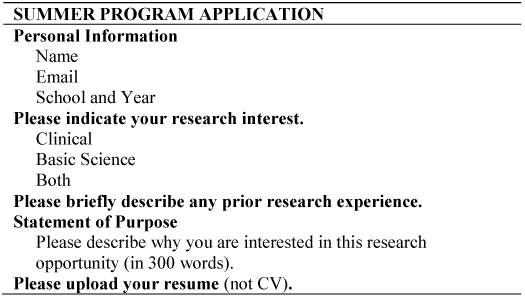
Application requirements and questions

**Figure 2.**
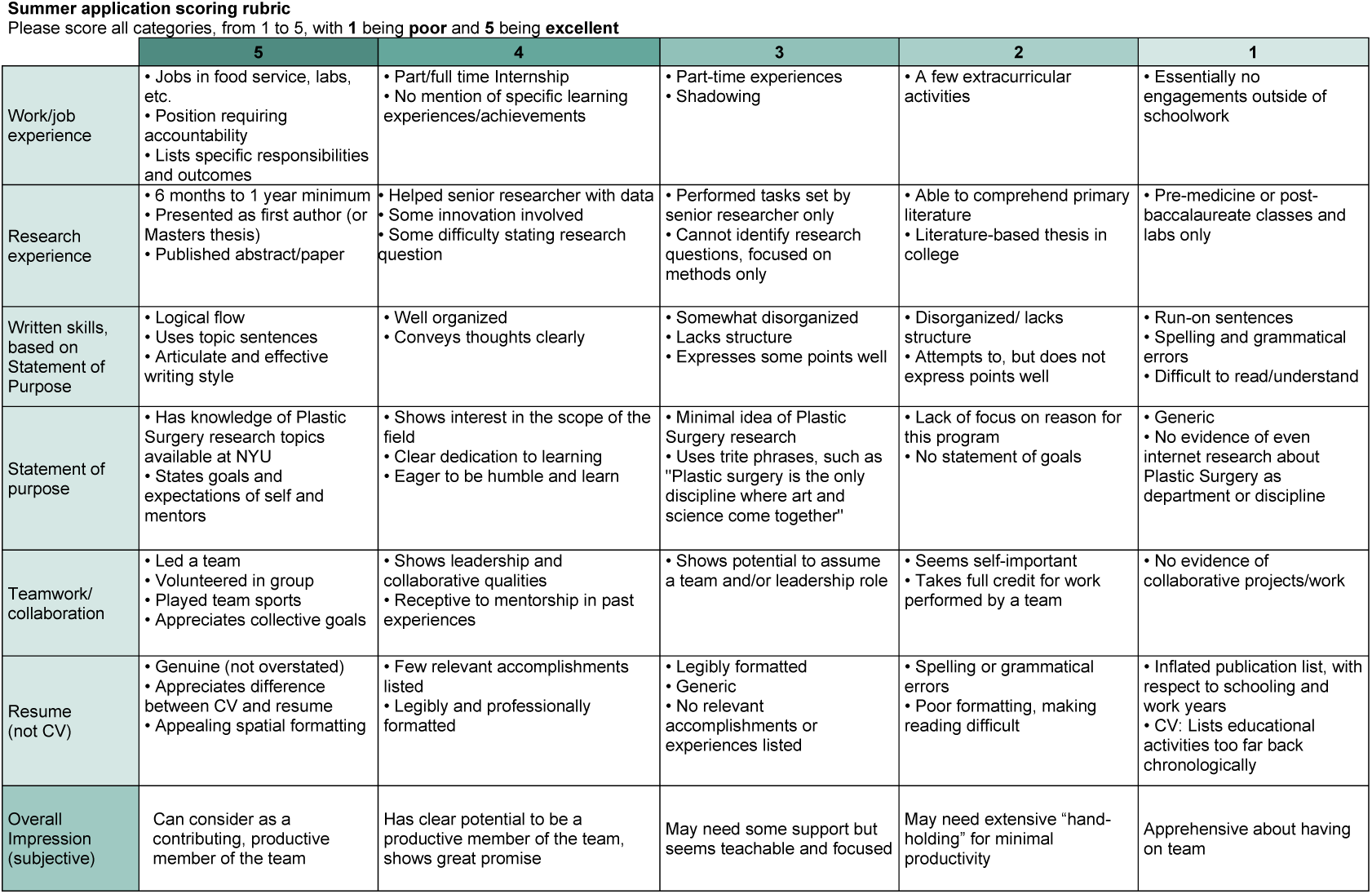
Scoring rubric

For each application, readers assign scores of one (lowest) through five (highest) per criteria based on the rubric, then assign an overall impression score out of five. During a meeting between all evaluators, overall impression scores are combined to generate a composite score out of ten. If impression scores are found to be discrepant, a group discussion is conducted to reach consensus and designate a composite score. Each member of the selection committee has an equal vote. In the data presented, interrater reliability was assessed retroactively by determination of intraclass correlation coefficient (ICC).^17^

In-person or web-based ten-minute interviews are granted to up to twenty applicants, depending on the application year and applicant pool. Interviews provide an opportunity to interact with prospective students and assess communication skills, curiosity about plastic surgery research topics, prior experience working within a team, and compatibility with current researchers. All application evaluators are present for the interviews and may refer to each interviewee’s application, resume and composite score. We then rank all interviewees and sub-group based on their expressed interest in clinical research, basic research, or both. We offer positions to the top three students per group and generate one combined waitlist in the event an applicant declines the offer. The timeline from application submission deadline to acceptance notification typically spans one month.

### Program Evaluation

In the last week of the program, direct mentors conduct individual exit interviews with each participant. We provide tailored feedback for each student and explain future research opportunities and recommendations for research years either as medical students or at a post-graduate level. Within one week of program conclusion, we distribute an online feedback form (**Table 2**) that rates the program on multiple parameters using a 5-point Likert scale. The responses are pooled by the director of the program, anonymized, and shared with the program team. On the administrative side, we take into account the feedback from students and staff to refine timeframes for administrative tasks, and discuss funds and operational dates for the following year. On the mentoring side, we conduct a de-orientation based on responses and discuss which feedback to incorporate into the following year’s program. Individual mentors receive feedback as well. We save all proposed changes for the following year in written form.

**Table 2.**
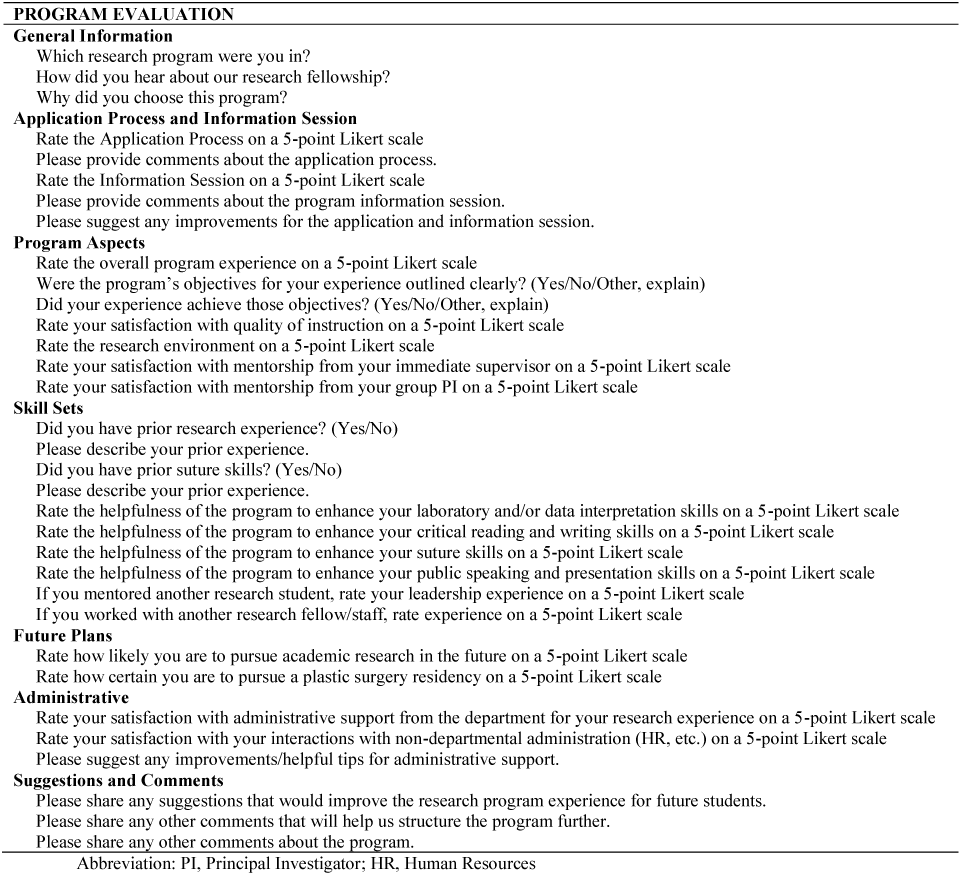
Program evaluation

Long-term program evaluation included analysis of continued research commitment and productivity. We defined research commitment as students who continued to perform research following program completion and/or returned to the department for dedicated time as a research fellow outside of the medical school curriculum. Productivity is measured by number of peer-reviewed articles published and searchable on PubMed following program completion and prior to medical school graduation. This is calculated as mean ± standard deviation. Figures were created using GraphPad Prism 8.0.2 (GraphPad Software, La Jolla, CA).

## RESULTS

### Applicant and Participant Composition

In 2013, this program started with seven students. When applications were instituted in 2014, 14 students applied and five participants enrolled. Program capacity has remained stable between three and eight participants per year. Over time, this has resulted in an increasingly selective acceptance rate starting at 36% in 2014 and reaching 7% in 2020. **Figure 3** depicts the composition of applicants from our institution compared to outside institutions, and shows the relative increase in program popularity over the last three years. This timeline aligns with more recent efforts to formalize applications and programming. **Figure 4** shows the composition of participants, which has included more participants from outside institutions in recent years, reflecting the growth in the applicant pool.

**Figure 3.**
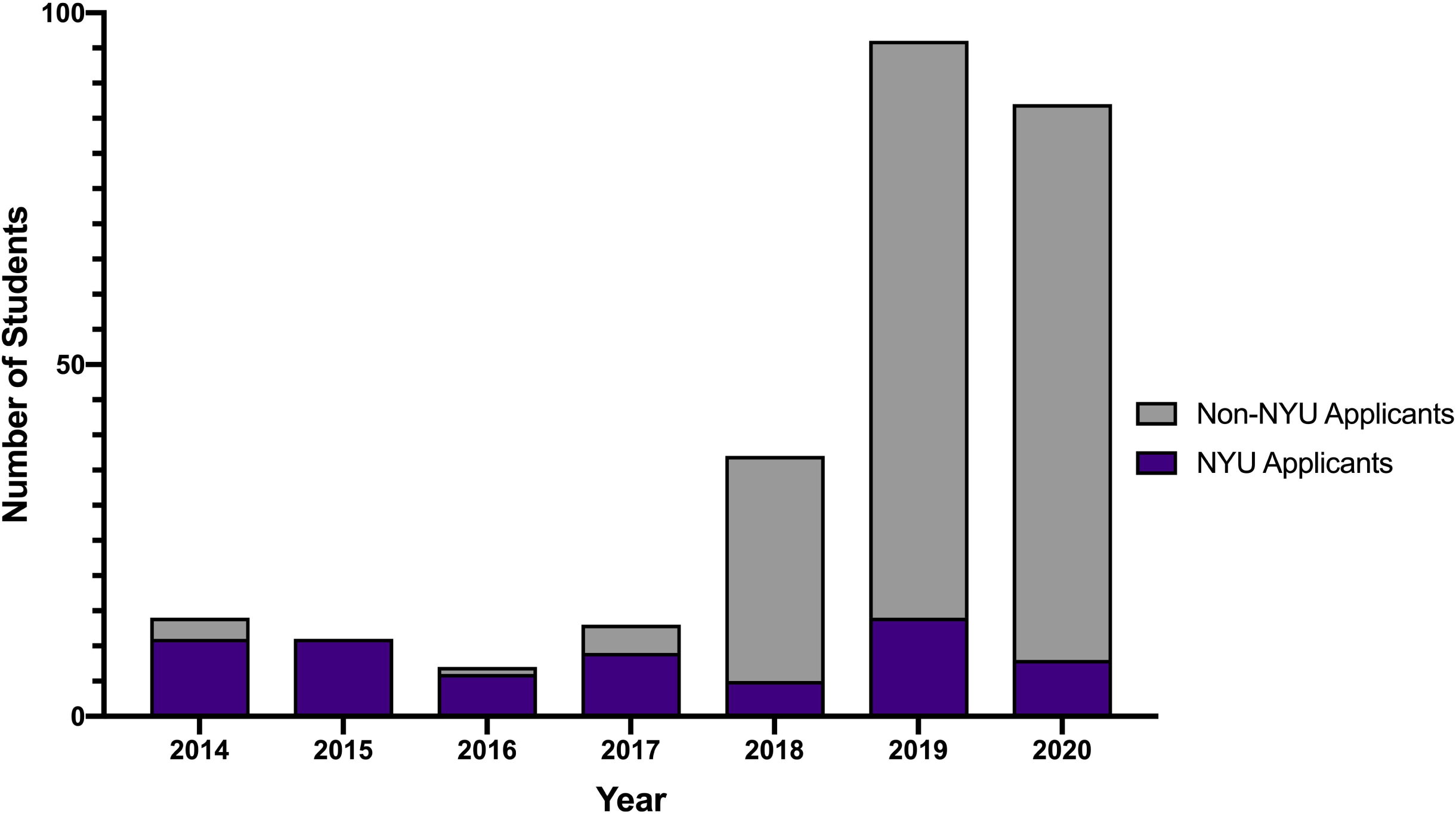
Applicant composition

**Figure 4.**
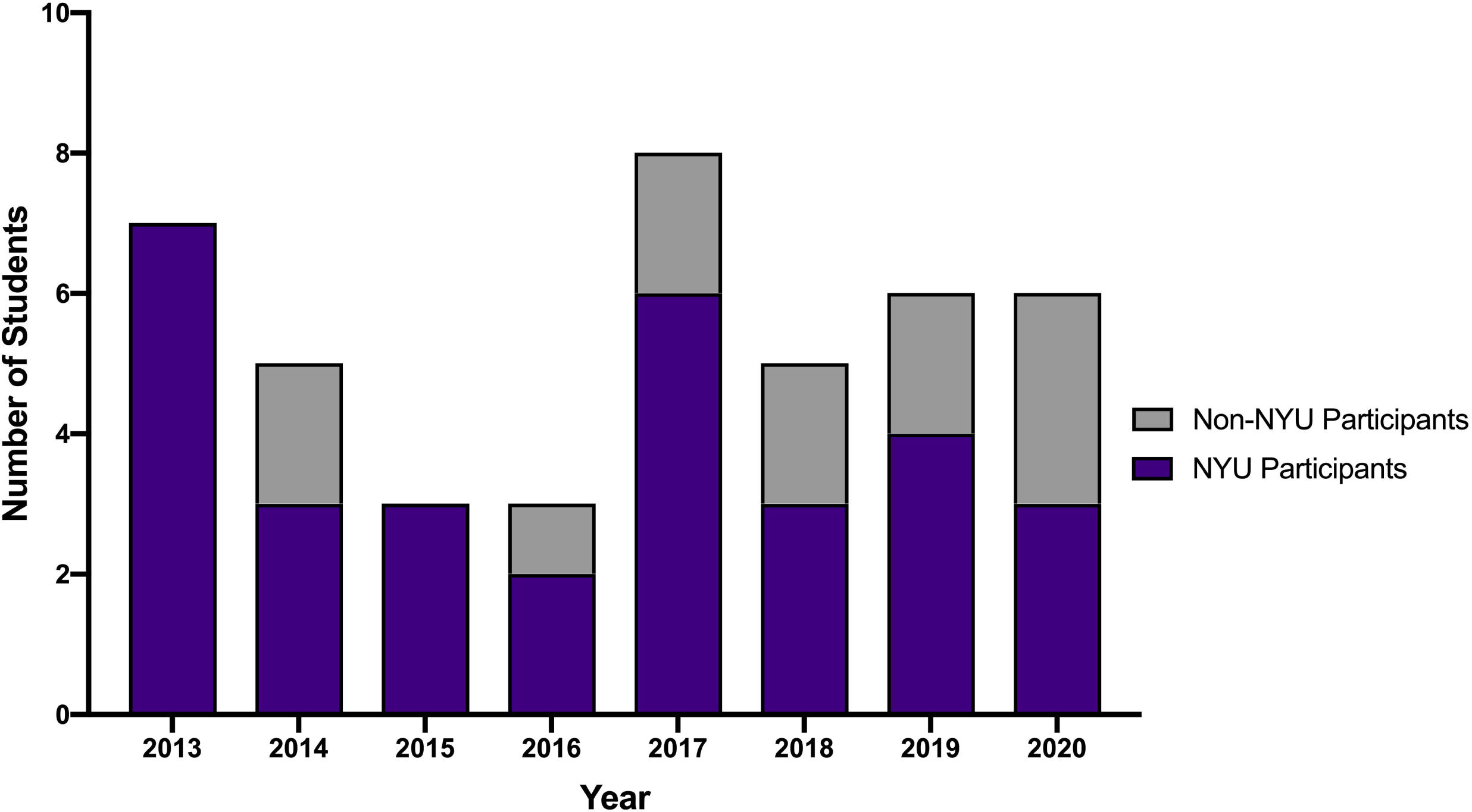
Participant composition

#### Reliable selection rubric

Of 87 students evaluated, 60 had two impression scores documented. Reasons for exclusion were those who did not qualify for the program, such as international medical school applicants, students beyond their first year of medical school, or those with a documented score only available from one evaluator due to the retrospective nature of the analysis. There was good to excellent interrater reliability between evaluators (ICC, 0.75; 95% confidence interval 0.59–0.85; p<0.001).

### Program Evaluation and Outcomes

Response rates for program evaluation questionnaires was 68% overall and 100% in the two most recent available surveys. Questionnaire responses for 2018 were not available as evaluations were not distributed that year. **Figure 5** depicts the trends in student satisfaction and rated influence of program enhancement of skills on a 5-point Likert scale (1, poor/unsatisfied; 5, excellent/extremely satisfied). Improvements in public speaking and presentation skills, and suture and clinical skills coincide with additional formal programming of mandatory departmental research presentations and structuring of suturing clinics, respectively. **Table 3** details the curricular evolution and **Figure 6** highlights the timeline of key curricular changes.

**Table 3.**
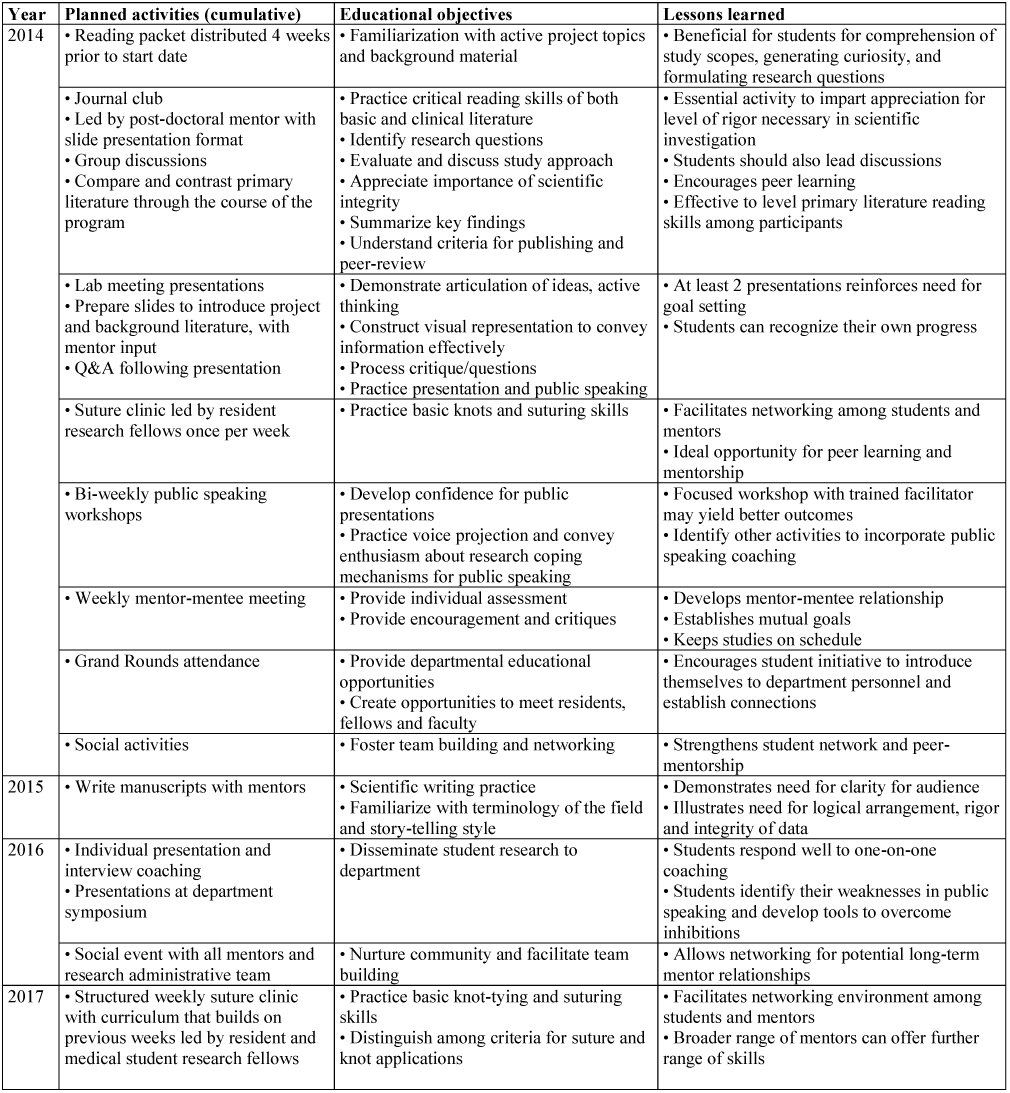
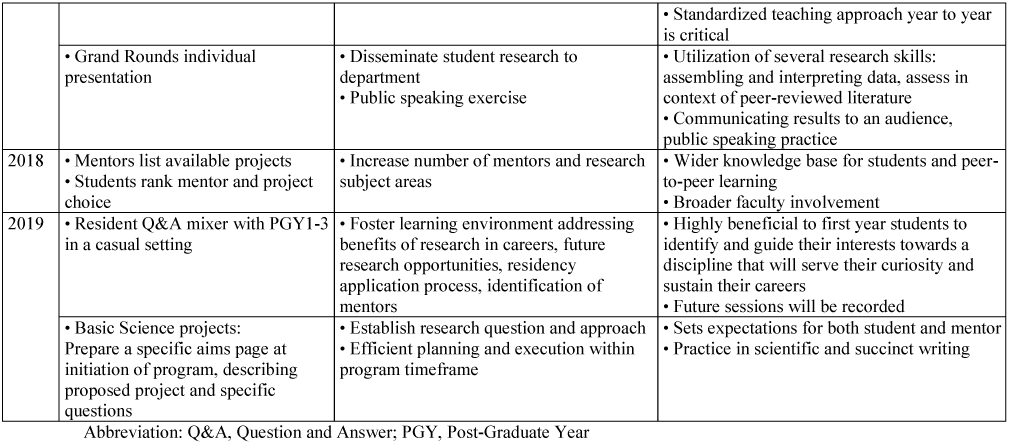
Evolution of curricular objectives and activities built upon yearly lessons learned *Legend:* This is a detailed, chronological evolution of the summer research program. Planned activities are cumulative across years and build upon the previous year’s lessons learned.

**Figure 5.**
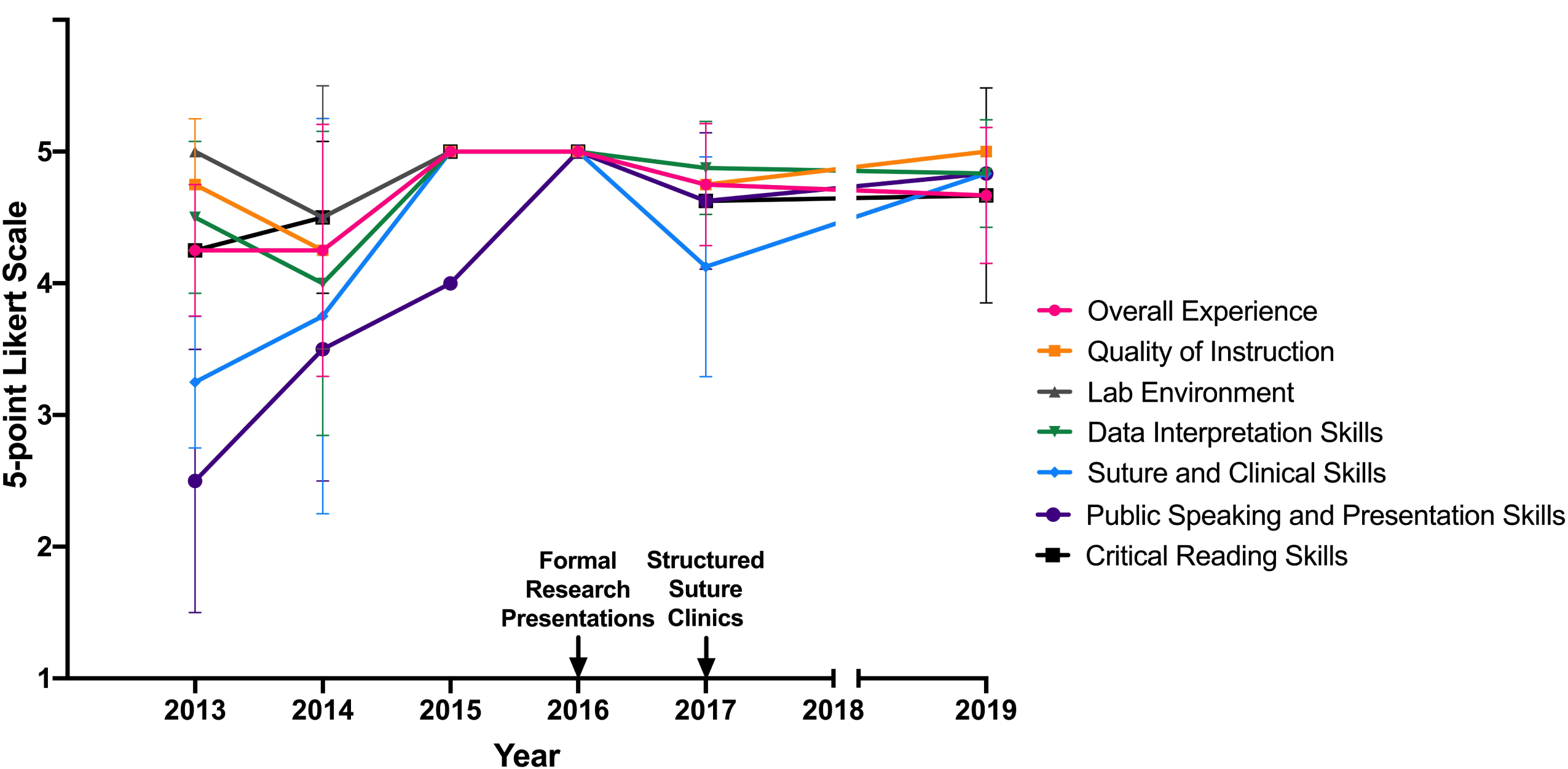
Program evaluation by students completing the summer research program

**Figure 6.**
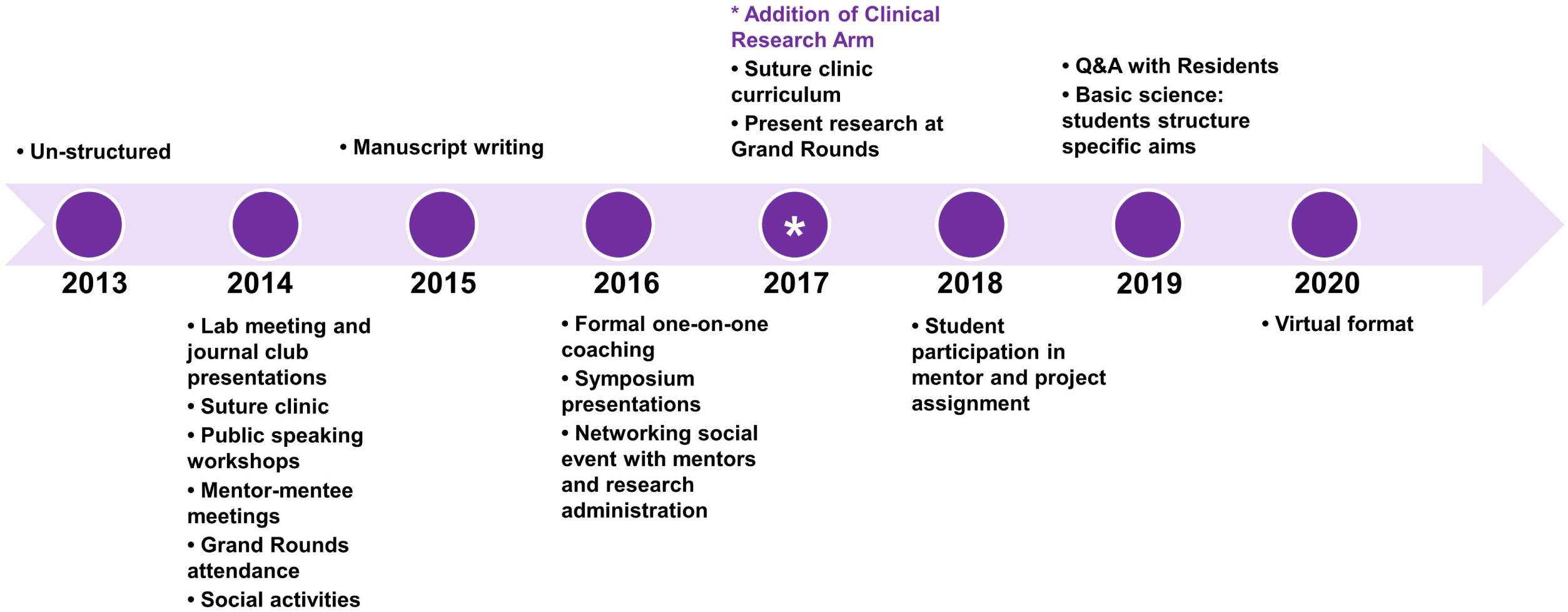
Timeline of key curricular developments

### Continuation of research productivity

On average per year from 2013 to 2019, 28% of program participants continued to conduct research with the plastic surgery department beyond the official program end date. From the cohorts of the summers spanning from 2013 to 2018, 29% of program participants returned to the department for a period of dedicated research outside of the medical school curriculum. The average number of publications per participant produced by the end of medical school was 6.9±4.0. Of these publications, 6.5±4.2 were affiliated with New York University and 5.0±4.2 were within the field of plastic surgery.

### Successful residency match

Amongst the participants who have now completed medical school (n=22), 95.5% successfully matched into a residency program on first attempt. Of these participants, 62% (n=13) matched into a surgical residency program with 54% (n=7) of these being an integrated plastic surgery residency.

## DISCUSSION

The medical student summer research program developed by our Department of Plastic Surgery is built around the institution’s academic missions of clinical excellence, education, and commitment to research advancement.^18^ Recognizing the importance of early career mentorship and initiation into evidence-based medicine and scientific investigative methods, the program was designed to provide students with foundational tools for ongoing self-development. Initially intended as an introduction to the field of plastic surgery through a research lens, the program has evolved into an incubator for highly dedicated and talented students who have consistently gone on to become exceptionally productive individuals and burgeoning leaders within their communities, as evidenced by their academic productivity.

Evidence-based decision making is an integral part of modern medicine, and responsible production, dissemination, and use of scientific evidence is crucial to patient safety and quality of surgical care.^19^ These elements are of paramount importance in the fast-paced innovative field of plastic surgery.^20^ While medical school curricula have seen substantial reconfigurations in recent years, opportunities for medical students to engage in applied clinical or basic science research within surgical departments are scarce, despite it being cited as an important factor in evaluating medical students for residency candidacy.^21,22^ Many medical schools lack affiliation with academic medical centers that have plastic surgery departments hosting residency training programs. When these do exist, the presence of a research infrastructure able to adequately accommodate medical students is rare.^23^ Our experience shows that such an opportunity can be created for a diverse pool of highly competitive applicants across the United States, with reproducible, quantifiable outcomes and a growing educational imprint.

### Mentorship

In our experience, the formalization of a mentor-mentee dynamic, even within the confines of an eight-week program, allows students to understand expectations, clearly delineate their own goals, focus their efforts in order to complete projects and leverage team-based learning to drive their personal development while contributing to the development of their peers. These benefits have been described widely.^24-28^ With the implementation of this model, we have seen students consistently complete the program with first author publications in addition to active involvement in other projects. Residents and fellows have the fulfilling and rewarding experience of providing mentorship to their juniors. Participating students have demonstrated autonomy, leadership and inquisitive critical thinking exemplified by their curricular participation, particularly through their research presentation at the completion of the program and the manuscripts they go on to publish. The mentorship relationships built within the program have persisted to become an anchoring force in students’ subsequent path through medical school and beyond.

### Research Project Management

The program’s structure revolves around common goals and expectations, and an outcomes-driven, personalized system for evaluating progress and discussing achievements and challenges. Participating students join a rigorous team of research fellows and faculty. This offers an immersive experience, with tailored feedback, team discussions and progress updates where the student can elaborate on their thinking process, discuss alternative approaches, and present data. With the team leaders’ supervision and guidance, students also collaborate to advance one another’s projects and mentor each other with regards to particular skills or areas of expertise developed throughout the course of their respective projects. By program completion, every student has typically been involved in literature review, study design and institutional review board (IRB) protocol development, data collection, analysis, interpretation, table and figure design and manuscript writing. This team-based approach allows engagement in the spectrum of activities that make up a successful research study. For students on the clinical research track, successful completion of the program entails completion of a first draft of a manuscript and oral presentation of their work at departmental grand rounds. For those on the basic science research track, oral presentation and abstract submission is typical.

### Educational Development

Successful participation and completion of research projects is marked by several milestones including abstract submission and presentation at conferences, submission and revision of manuscripts and grand rounds presentations. Students additionally develop their interpersonal and academic prowess by attending weekly didactic sessions alongside residents and faculty, and engage in weekly journal clubs, further developing their critical thinking, communication, and oral presentation skills. This is supported by the good to excellent rating by students of the helpfulness of the summer program at enhancing their skills in data interpretation, critical reading and writing, public speaking and presentation. Each student chooses a peer-reviewed article and presents to the group, with emphasis on critical appraisal and clinical correlation. Journal clubs are followed by hands-on surgical skills sessions. Students typically join the program with little to no exposure to surgical technique and complete the program with a basic understanding of instruments and materials and reproducible beginner-level suturing and knot-tying skills as represented by their program evaluation survey responses. When schedules coincide, students are invited to participate in resident cadaver dissection sessions, allowing them to strengthen their knowledge of anatomy and apply the skills they are learning to clinically-relevant scenarios.

### Limitations

The evolution of this program is due to our commitment to quality culture and enhancement. The changes observed therefore reflect the identified limitations and proposed solutions. This is a report of the evolution of a summer research program at a research institution in an urban setting with a Department of Plastic Surgery that can formally accommodate approximately six summer research students per year. Though we present the framework for developing your own summer research program, the information presented here should be considered within the context it was successful, and appropriately applied to new contexts. The limitations of this study include its retrospective nature that lends to missing survey data in 2018. This is not a controlled study and therefore the long-term outcomes measured cannot be solely attributed to participation in the program, nor is that the intent of the analysis. Data to support the use of the scoring rubric would benefit from continued analysis to collect a larger sample size over the years.

### Future Directions

Facing the challenges imposed by the evolving COVID-19 pandemic, we implemented remote-learning and collaborative telecommunication platforms to enable students to continue to benefit from the summer research program. Participant feedback will continue to play a vital role in the future success of the program. Future investigations will include assessment of the use of remote versus in-person platforms, including transition to virtual journal clubs, suture and knot-tying lessons, and group-based as well as individual feedback sessions. Ultimately, our hope is to expand the parameters of the program, potentially expanding the application pool to invite international candidates to participate. We anticipate that virtual sessions may have the potential to reach a wider audience and therefore may be adopted and incorporated into the syllabus even as in-person sessions resume.

## CONCLUSION

A research program designed for pre-clinical medical students interested in plastic and reconstructive surgery can reliably achieve academic and educational goals. It also provides students with mentorship opportunities, a professional network, and the tools for self-guided learning and subsequent career development.

## ACKNOWLEDGMENTS

We are grateful to Dr. Rami Kantar for his dedication as a mentor to this program, Ms. Aimee Chow, Mr. Hong Zheng and Ms. G. Leslie Bernstein for administrative coordination, and staff of the Department of Plastic Surgery for their support.

## Notes

**Financial disclosure statement:** The authors have no relevant disclosures to report.

### Competing Interest Statement

The authors have declared no competing interest.

